# Longitudinal three-photon imaging for tracking amyloid plaques and vascular degeneration in a mouse model of Alzheimer’s disease

**DOI:** 10.1101/2025.08.12.669914

**Authors:** Eline Stas, Mengke Yang, Simon Schultz, Mary Ann Go

**Author notes:** Co-First Author (Eline Stas, Mengke Yang).

## Abstract

**Significance:** Vascular abnormalities may contribute to amyloid-beta accumulation and neurotoxicity in Alzheimer’s disease (AD). Monitoring vascular degeneration as AD progresses is essential. Three-photon fluorescence microscopy (3PM) enables high-resolution deep tissue imaging with minimal invasiveness and photodamage.

**Aim:** This proof-of-concept study established a longitudinal 3P imaging pipeline to quantify vascular and amyloid plaque changes in the APP^NL-G-F^ mouse model.

**Approach:** A cranial window allowed repeated 3P imaging at four-week intervals beginning at five weeks after surgery. Vessels labelled with Texas-Red were segmented using DeepVess, while plaques labelled with methoxy-XO4 were segmented using custom scripts. Quantitative analyses assessed vascular parameters (diameter, tortuosity, length, inter-vessel distance, total volume) and plaque metrics (radius, total volume).

**Results:** We imaged the same field over 4 weeks quantifying an overall decrease in vasculature and increase in amyloid plaques between two sessions. Significant changes in vessel diameter, inter-vessel distance, as well as alterations in vessel length and plaques radius were observed. Changes in vessel tortuosity were not significant.

**Conclusions:** We demonstrate the potential of three-photon imaging to track vascular and amyloid-related changes in deep cortical structures. It offers a tool for studying the interplay between vascular and amyloid pathologies in AD, supporting future research into disease mechanisms and therapeutic strategies.

## 1 Introduction

Alzheimer’s disease (AD) is the leading cause of dementia worldwide, impacting the quality of human life and imposing significant healthcare costs.^1^ A widely accepted theory, the amyloid hypothesis, suggests that cognitive decline in AD patients is partly due to the accumulation of extracellular amyloid beta (A*β*) peptides in the brain.^2^ This accumulation is believed to trigger a cascade of pathological events, including the formation of intracellular neurofibrillary tangles composed of hyperphosphorylated tau and subsequent neuronal loss.^3, 4^ Nonetheless, while amyloid pathologies have been the primary focus, emerging studies highlight the vascular system as a major factor in the progression of the disease.^5^

To meet its high energy demands, the brain relies on a constant supply of oxygen and energy substrates. As the brain has no long-term energy storage, regional cerebral blood flow is precisely regulated to align with the local energy requirements of nervous tissue. Any disruption in this adaptive process or in the delivery of necessary substrates can lead to imbalances in homeostasis, tissue damage, and functional impairment.^6^ These vascular abnormalities have been shown to not only contribute to A*β* accumulation, but also exacerbate neurotoxicity in AD.^7–9^ It is believed that the interplay between vascular degeneration and A*β* deposition is central to understanding the pathophysiology of AD progression. Brains of human AD patients commonly exhibit distinct vascular anomalies, including significant reductions in vascular diameter,^10^ density,^11^ and blood flow,^12^ while vessel tortuosity and length tend to increase.^13, 14^ However, the precise mechanisms by which vascular abnormalities contribute to amyloid plaque buildup and subsequent neurotoxicity in AD remain unclear.^15^

Animal models are crucial for elucidating the pathophysiological pathways of AD and developing preventive, diagnostic, and therapeutic strategies. While wild-type mice do not form senile plaques or neurofibrillary tangles,^16^ genetically engineered mice consisting of familial AD (FAD)-linked mutations in genes like *APP, PSEN1*, and *PSEN2* are employed to mimic amyloidosis features of FAD-AD.^17^ Most AD cases are caused by sporadic AD, however no animal models mimicking this form have been developed yet. Early transgenic models overexpressed amyloid precursor protein (APP), leading to excessive A*β* production and accumulation of APP fragments.^18^ However, APP overexpression can introduce artifacts not representative of human pathology.

The APP^NL-G-F^ knock-in mouse model, which avoids APP overexpression by introducing specific mutations into the endogenous *APP* gene, provides a more accurate representation of preclinical AD.^19, 20^ This model incorporates the Swedish (NL), Arctic (G), and Iberian (F) mutations, with homozygous mice exhibiting greater A*β* accumulation starting at 2 months than heterozygous ones starting at 4 months.^21, 22^ It is considered a standard for studying mechanisms of A*β* amyloidosis.^18^ FAD typically manifests in humans in the heterozygous state, therefore the heterozygous APP^NL-G-F^ mutation more accurately models human pathology.^23^ Although other AD mouse models such as APP/PS1, 5xFAD, and APP23 have been utilized to study changes in vascular morphology and plaque progression,^24–27^ chronic imaging of vascular and amyloid interactions in the APP^NL-G-F^ knock-in model has not been thoroughly investigated. This gap presents an opportunity to examine the relationship between vascular abnormalities and amyloid plaque formation in a model that more closely replicates human A*β* pathology.

Despite advancements in AD research, the disease’s complex pathogenesis remains poorly understood, highlighting the necessity for advanced longitudinal imaging methods to sequentially visualize brain structures and gain further insights into the mechanisms of the disease. Advanced imaging techniques are essential for studying the dynamics of vascular degeneration and amyloidpathology *in vivo*. Three-photon fluorescence microscopy (3PM) has significantly extended the penetration depth of high-resolution optical imaging, enabling the visualization of individual cells in preclinical rodent models with high spatial and temporal resolution.^28^ Compared to traditional noninvasive imaging techniques such as CT, MRI, and PET, 3PM provides cellular-level resolution at the cost of requiring surgical cranial access and the use of high-power laser excitation. Nevertheless, when compared with other optical microscopy approaches, 3PM achieves deep-tissue imaging with relatively minimal photodamage.^29–32^ Additionally, 3PM employs pulsed laser sources that deliver high peak power for excitation at low repetition rates, resulting in low phototoxicity.^33^

Specifically, 3PM enables deep tissue imaging using long excitation wavelengths (1300–1700 nm) and low average power, minimizing photodamage and phototoxicity.^33, 34^ This approach reduces out-of-focus signal and enhances the signal-to-background ratio (SBR), allowing imaging at depths up to 1.4 mm—surpassing the limitations of two-photon fluorescence microscopy (2PM), which is often restricted to depths of 700 *µ*m.^31, 35^ Multicolor 3PM, using fluorescent dyes and genetically encoded markers, has successfully imaged structures such as GCaMP6s-labeled neurons, Texas Red–labeled blood vessels, and third-harmonic generation (THG) signals from myelinated axons up to 1.1 mm deep. THG is a nonlinear optical phenomenon that enables label-free brain imaging by generating a signal at three times the original laser frequency, with its efficiency enhanced by structural discontinuities within the focal volume.^36^ However, simultaneous multicolour imaging of vasculature and A*β* using 3PM has not been demonstrated, and longitudinal imaging using 2PM has been limited to superficial cortical regions within layer five.^30, 37^

Our study aims to address this gap by establishing a longitudinal imaging methodology for deep cortical structures in the brain, utilizing a 3PM imaging pipeline to investigate vascular and amyloid pathologies in the APP^NL-G-F^ knock-in mouse model. By leveraging the deep imaging capabilities of 3PM, we seek to enable future studies to explore the dynamics of vascular degeneration and A*β* plaque accumulation. This approach will provide more accurate and translatable insights into the physiological mechanisms underlying AD progression, ultimately supporting the development of effective preventive and therapeutic strategies. Understanding these mechanisms could significantly impact future research and clinical practices in AD, paving the way for improved diagnostic and therapeutic approaches.

## 2 Methods

### 2.1 Three-Photon Microscope Setup

The 3P microscope [Fig. 1(a)] consisted of a 40 W fibre laser (Satsuma), an optical parametric amplifier (OPA, Mango) running at a repetition rate of 1 MHz, and a hyperscope (Scientifica). The laser pulses, with a width of 65 fs at 1340 nm, passed through a pulse compressor composed of mirrors (M3, M4, M5) and prisms (P1, P2). The beam was then directed through a series of mirrors (M1-M15), a half-wave plate and a beam-splitter for beam conditioning. The final power under the objective ranged between 165–220 mW. A tiltable objective mount was used, supporting a 25× water-immersion objective (XLPlan NA=1.0, Olympus), with an ultrasound gel as the immersion liquid. The signals were divided into two detection channels using two photomultiplier tubes (PMTs) and dichroic beam splitters with specific wavelength filters (ET440/40 and ET620/60 from Thorlabs). The ET440/40 filter was used to detect amyloid plaques via methoxy-XO4 fluorescence,^38^ while the ET620/60 filter was used to detect vessels via Texas-Red fluorescence.^36^ Before reaching the objective lens, a beam expander was used to ensure the filling of the back aperture for optimal imaging resolution. The focal spot sizes along the x, y, and z axes were determined using 0.20-*µ*m-diameter fluorescent beads, calculated as the full-width at half-maximum (FWHM) [Fig. 1b]. Lateral resolution was measured to be 0.87 ± 0.12 *µ*m while axial resolution was 4.59 ± 0.48 *µ*m (mean ± s.e.m, n=5).

**Fig 1.**
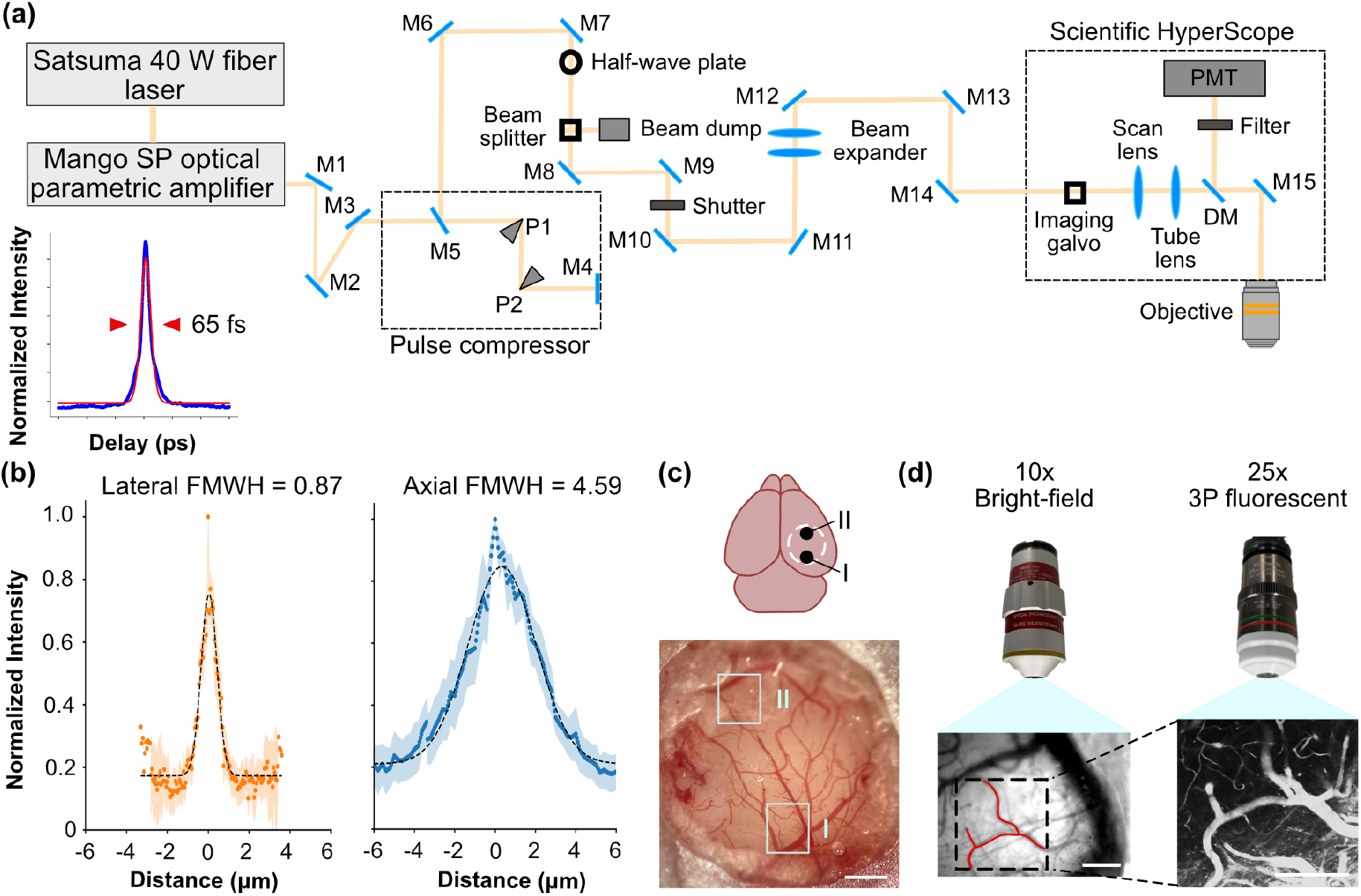
Imaging pipeline using a three-photon fluorescent microscope. (a) Schematic of the three-photon excitation imaging system with a pulse width of 65 fs. The system uses a Satsuma 40 W fiber laser and Mango SP optical parametric amplifier as the excitation sources. Key components include mirrors (M1-M15), a prism pulse compressor (P1-P2), a half-wave plate, a beam splitter, a beam expander, a shutter, a dichroic mirror (DM), scan and tube lenses, and an objective lens. The fluorescence signal is collected using a photomultiplier tube (PMT). (*Lower left*) Measured pulse width shown in blue with sech^2^ fit (pulse width 65 fs) shown in red. (b) Intensity profile in the lateral (orange) and axial (blue) distance of 200-nm fluorescent beads (n=5). (c) Cranial window preparation and imaging locations: Diagram and bright-field image of a mouse brain with a cranial window. Imaging FOVs (I: ML: 2.25 mm, AP: -3 mm and II: ML: 2.25 mm, AP: -1 mm) are marked. Scale bar: 1 mm. (d) Imaging procedure and objectives: Bright-field and three-photon fluorescent images are acquired using 10x and 25x objectives to create vascular maps. Scale bars: 200 µm.

### 2.2 Animals

All experimental procedures complied with the project license (PPL number: PP6988384), Home Office, and Imperial College London institutional norms. Heterozygous APP^NL-G-F^ knock-in mice^19^ with one copy of the mutant APP gene were employed as AD models. The humanized A*β* buildup induced by mutations results in similar disease development, making the mouse model suited for studying the link between amyloid plaques and blood vessels. Six mice were used: one female aged 8.9-to 9.9-months and two females aged 6.5- to 7.5- months for chronic imaging, and a 10- and 11-month-old mouse for image analysis pipeline development; a 1-month-old wildtype (C57BL/6) mouse was also imaged for pipeline development.

### 2.3 Cortical Surgery and Viral Injection

Mice were given Carprofen (5 mg/kg) and buprenorphine (0.07 mg/kg) for pre-operative pain management and were operated on under 1.5–3% isoflurane anesthesia. A heated blanket and rectal thermometer were used to maintain body temperature at 37°C. A circular craniotomy was made over two fields of view (FOV) in the cortex. FOV I was located 2.25 mm ML, -3 mm AP from bregma and FOV II 2.25 mm ML, -1 AP mm from bregma [Fig. 1c]. The craniotomy was outlined with a 3 mm biopsy punch, and the skull piece removed using a dental drill. A 3 mm coverslip was attached to a 5 mm coverslip using optical glue and attached on to the craniotomy together with a head plate using dental cement. After surgery, a photograph was taken as a reference for vascular mapping. Mice were allowed to heal for 35 days before beginning imaging.^39, 40^ For one mouse imaged for pipeline development, 5uL of the virus CAG-NLS-GFP (Addgene 104061, titer 2.5 × 10^13^ GC/ml) was injected through the facial vein at P0.^41^ The virus contains a green genetically encoded nuclear fluorescent protein. Imaging was done at P32.

### 2.4 Chronic Imaging

Mice underwent two imaging sessions with a four-week interval. Mice were anesthetized with 1–1.2% isoflurane, maintained at 37°C, and monitored via an infrared camera. Amyloid plaques were visualized using methoxy-X04 (10% DMSO, 45% propylene glycol, 45% saline), injected intraperitoneally 24 hours before imaging. Vessels were imaged with Texas-Red (50 mg/kg at week 5, 25 mg/kg at week 9), delivered via tail-vein injection 30 minutes before *in vivo* imaging. Using a stereomicroscope before imaging, vessel patterns were compared to the vascular images taken immediately after surgery. Vessels exhibiting distinct morphological features such as large diameter, pronounced curvature, bifurcations, or crossings were identified and used as landmarks. Then the mouse was stabilized under the microscope by fixing it in a holder using using the head plate and these characteristic features were referenced during both imaging sessions to accurately locate the same region using a 10 × air-immersion objective (TL 10X-2P, Thorlabs). Once the area was located, it was imaged with a 25× water-immersion objective (XL Plan NA=1.0, Olympus) [Fig. 1(e)]. During chronic imaging, the power under the objective ranged from 165-182 mW with inter-session power variability for each FOV not exceeding 8%. To assess measurement stability, a neonatal mouse lung slice was imaged at different power levels, and the width of several structures were measured. Size variability was minimal across an 8% power difference and did not exceed 6% [Fig. S1]. Image stacks (512×512 pixels, 500×500*µ*m FOV, 5 *µ*m step size along the z-axis) were taken with 5–7 frames per plane, depending on image quality, using ScanImage.^42^

### 2.5 Image Processing

Before processing the images, stacks were manually inspected for obvious signs of signal loss between sessions. Two datasets were excluded from the analysis: Mouse 2 FOV I showed vascular regrowth in Session 2 rendering longitudinal comparison with Session 1 invalid due to partial loss of signal in the FOV, and Mouse 3 FOV II exhibited insufficient SNR in both sessions. Images from selected stacks were then intensity-matched using adaptive histogram equalization, and crosstalk was removed. Preprocessing included dura removal, normalization, noise reduction^25^ [Fig. 2(a)]. Motion artifact was assessed by binarizing the raw data and selecting several single-vascular structures at different depths in each volume, and calculating the shift of the center position of the vascular structure signal brightness of each frame compared to the previous frame [Fig. S2]. The shift was found to be 2 pixels at maximum, therefore motion correction was not applied. The pipeline was run using MATLAB R2023b and Python.

**Fig 2.**
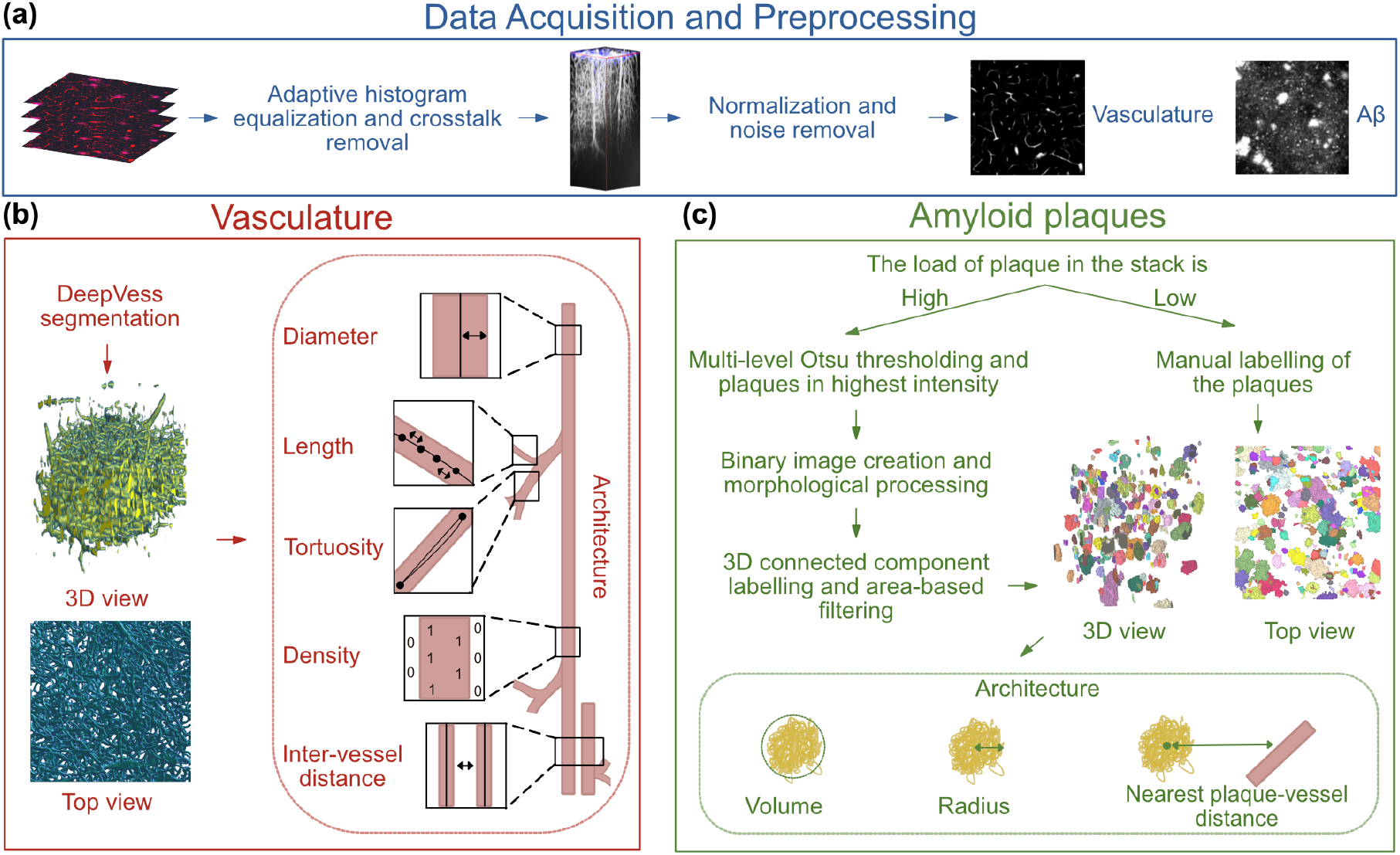
Segmentation and quantification pipeline for blood vessels and amyloid plaques in multiphoton images. (a) Data Acquisition and Preprocessing: adaptive histogram equalization, crosstalk removal, normalization, noise. (b) Vasculature Analysis: segmentation using the DeepVess pipeline, generating 3D and top-view visualizations. Quantification includes measures of (i) diameter, (ii) length, (iii) tortuosity, (iv) density, and (v) inter-vessel distance. (c) Amyloid Plaque Analysis: segmentation through multi-level Otsu thresholding for low plaque load or manual labeling for high plaque load. Quantification includes (i) plaque volume, (ii) plaque radius, and (iii) the nearest plaque-vessel distance.

#### 2.5.1 Blood Vessels

Vessels were segmented using the DeepVess 3D Convolutional Neural Network (CNN) model,^25^ which consists of six 3D convolutional layers with ReLU activation and max-pooling, followed by a softmax layer for classification. Dice similarity coefficients were computed for each stack by manually annotating six slices spaced 100 µm apart to provide representative validation across cortical depth. When the Dice coefficient was below 0.7, the corresponding stack was manually corrected using Napari^43^ to improve segmentation accuracy. The Dice coefficients are reported in Table 2.1.

After segmentation, vessel parameters, including diameter, length, tortuosity, inter-vessel distance, and density, were calculated based on previous works^44–47^ [Fig. 2(b)]. The diameter *D* of a vessel segment was determined as twice the median radius measured along the vessel’s centerline, where the radius *r*_*j*_ at each point was the minimum distance to the nearest background voxel:

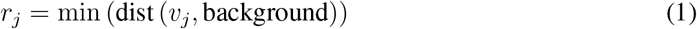

**Table 1.** Dice similarity coefficients (DeepVess vs. manual) by mouse, field of view (FOV), and session. A QC threshold of 0.70 was used to flag cases for manual correction.

| Mouse | FOV | Session | Dice | QC action |
| --- | --- | --- | --- | --- |
| Mouse 1 | I | Session 1 | 0.83 | Accepted |
| Mouse 1 | I | Session 2 | 0.70 | Accepted |
| Mouse 1 | II | Session 1 | 0.80 | Accepted |
| Mouse 1 | II | Session 2 | 0.74 | Accepted |
| Mouse 2 | II | Session 1 | 0.79 | Accepted |
| Mouse 2 | II | Session 2 | 0.72 | Accepted |
| Mouse 3 | I | Session 1 | 0.53 | Manual correction |
| Mouse 3 | I | Session 2 | 0.63 | Manual correction |

Here, *v*_*j*_ is the voxel at the *j*^*th*^ point along the vessel. The diameter *D* is then given by:

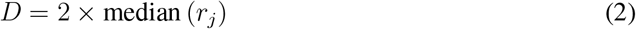

The length *L* of a vessel segment was calculated by summing the Euclidean distances between consecutive points along the vessel centerline:

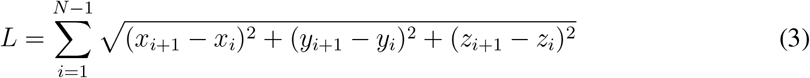

where (*x*_*i*_, *y*_*i*_, *z*_*i*_) are the coordinates of the *i*^*th*^ point along the vessel, and *N* is the total number of points. Tortuosity *T* was defined as the ratio of the vessel segment’s path length *L* to the straight-line distance between its endpoints:

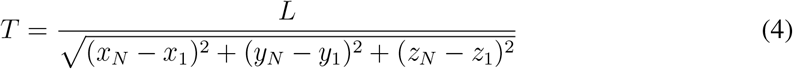

Here, (*x*_1_, *y*_1_, *z*_1_) and (*x*_*N*_, *y*_*N*_, *z*_*N*_) represent the coordinates of the starting and ending points of the vessel segment.

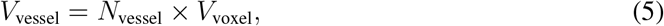

This physical volume *V*_voxel_ is given by the product of the in-plane pixel size and axial spacing:

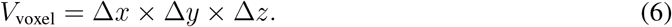

Finally, the inter-vessel distance (IVD) was calculated using a single midpoint per vessel segment. Specifically, for each skeletonized segment we took the centroid (mean of its [x,y,z] skeleton coordinates). We then measured Euclidean distances from that centroid to the centroids of all other segments and reported the minimum of these distances for that segment:

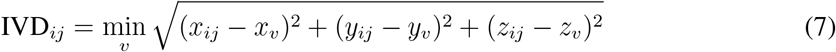

where (*x*_*ij*_, *y*_*ij*_, *z*_*ij*_) are the coordinates of the midpoint of vessel segment *ij*, and (*x*_*v*_, *y*_*v*_, *z*_*v*_) are the coordinates of the nearest segment midpoint *v*.

### 2.6 Amyloid Plaques

A Python script was developed to segment Amyloid plaques either semi-automatically using the SciPy package^48^ with manual correction via Napari,^43^ or fully manually [Fig. 2(c)]. Semi-automatic segmentation was used when the data showed more than 45 plaques with good imaging quality; otherwise, manual annotation was performed. For semi-automatic segmentation, four intensity thresholds were generated using multi-level Otsu thresholding, with plaques identified as regions with the two highest intensities. A binary image was then created, and morphological operations filled small holes and removed small objects. Area-based filtering was applied after 3D-connected component labeling. Regions smaller than 49 *µ*m^2^ were excluded.^49^ The 500 × 500 labeled areas in 2D planes (XY) with a 0.98 *µ*m pixel size were summed and transformed into 3D using a 5 *µ*m (Z) step size. The volume of each plaque was calculated directly from the 2D segmentations by summing per-slice areas and multiplying by the slice thickness. Using that volume, a radius was computed as the equivalent-sphere radius from this measured volume for comparison purposes.

### 2.7 Nearest Plaque-to-Vessel Distance

The coordinates of plaque centroids were retrieved from the segmented 3D binary data set. The closest neighbouring vessel of each centroid was identified using 3D Euclidean distances analysis.^50^

### 2.8 Data Analysis

Python was used for statistical analysis using the SciPy^51^ and Scikit-learn^52^ packages. The signal-to-background ratio (SBR) of imaging stacks was determined as the average of the brightest 15 and darkest 15 pixels in each profile.^53^ The signal strength was quantified by averaging the intensity of the top 0.5% of pixels in each frame.^32^ Data was tested for normality before statistical testing of vascular parameters and amyloid plaques between imaging sessions within a mouse. Mann-Whitney *U* tests were used as statistical testing. We reported distribution means as sample mean ± standard error of the mean unless specified otherwise.

## 3 Results

### 3.1 3PM Imaging: Dual-Channel Visualization of Vessels with Amyloid Plaques or Neurons

Using 1340-nm three-photon (3P) excitation, we acquired deep-tissue fluorescence images of the mouse cortex labeled with multiple fluorescent probes [Fig. 3]. In one configuration, vessels were visualized by Texas Red (magenta), while amyloid plaques were selectively stained with methoxy-XO4 (cyan). An additional signal from third-harmonic generation (THG, shown in yellow) shows the dura matter. In a second configuration, Texas Red (magenta) was used to label the vasculature, while neurons labeled with GFP (green) highlighted neuronal structure. The resulting 3D renderings captured both vascular architecture and either plaque or neuronal features in a single scan, underscoring the utility of 3PM for simultaneous, deep-tissue imaging of multiple tissue components.

**Fig 3.**
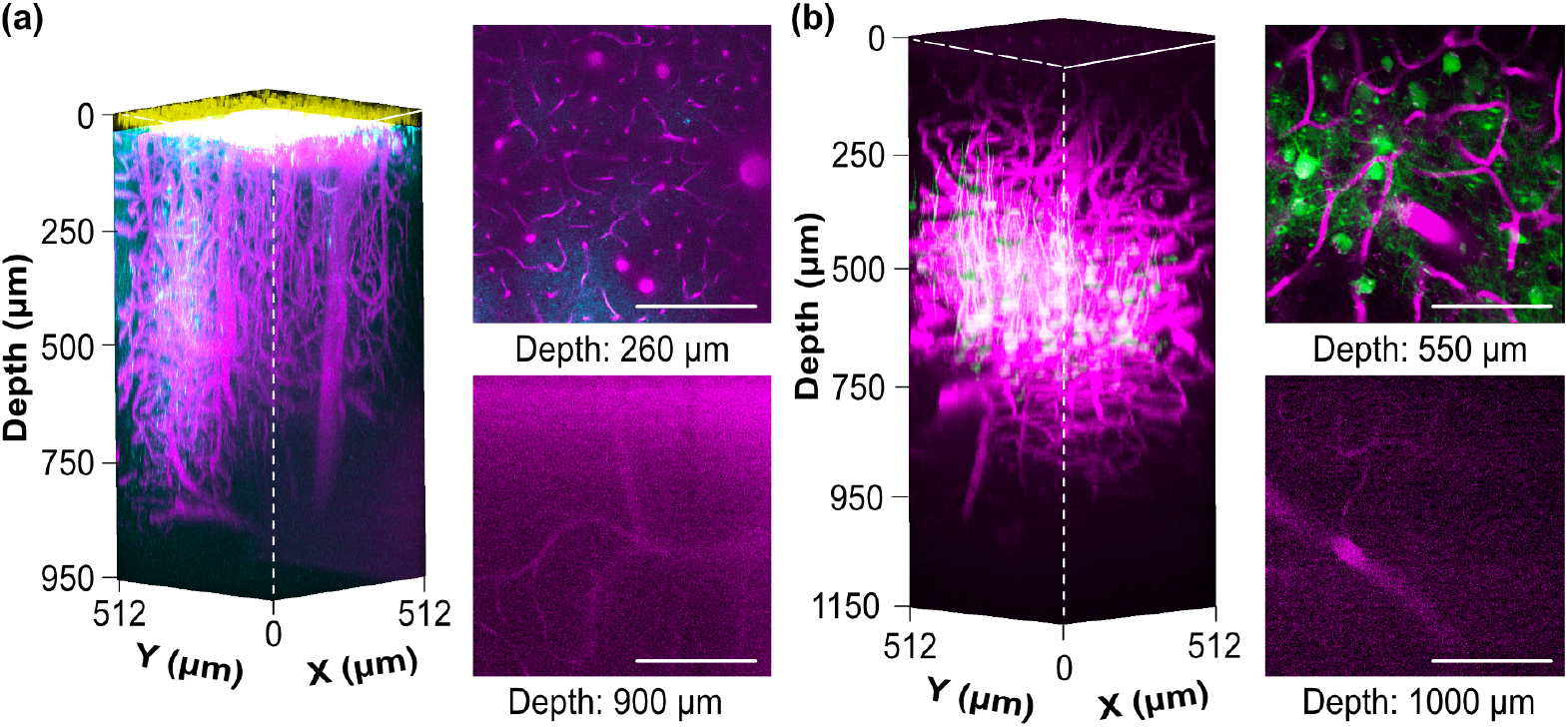
Multicolor 3P fluorescence images using 1340-nm excitation. 3D reconstruction of (a) vessels dyed with Texas Red (magenta), plaques with methoxy-XO4 (cyan), and THG (yellow) image stacks taken with 5 *µ*m z-steps, and (b) vessels dyed with Texas Red (magenta) and neurons labeled with GFP (green) taken with 10 µm z-steps. Stacks are normalized by adjusting the intensity of each slice based on the intensity range of the last non-overexposed slice (95% of maximum brightness) to ensure consistent brightness across the stack. The maximum laser power was (a) 182 mW and (b) 200 mW under the objective lens. Scale bars: 200 *µ*m.

### 3.2 Consistent Field of View Tracking Across Two Imaging Sessions

We conducted longitudinal imaging of the same FOV in APP^NL-G-F^ knock-in mice at weeks 5 and 9 after surgery to assess the progression of vascular and amyloid pathologies. Using vascular landmarks identified in postoperative bright-field images [Fig. 4(a)], we localized the FOV with a 10× air-immersion objective. High-resolution imaging was performed with a 25× water-immersion objective, enabling 3PM of cortical depths ranging from 45 to 795 *µ*m.

**Fig 4.**
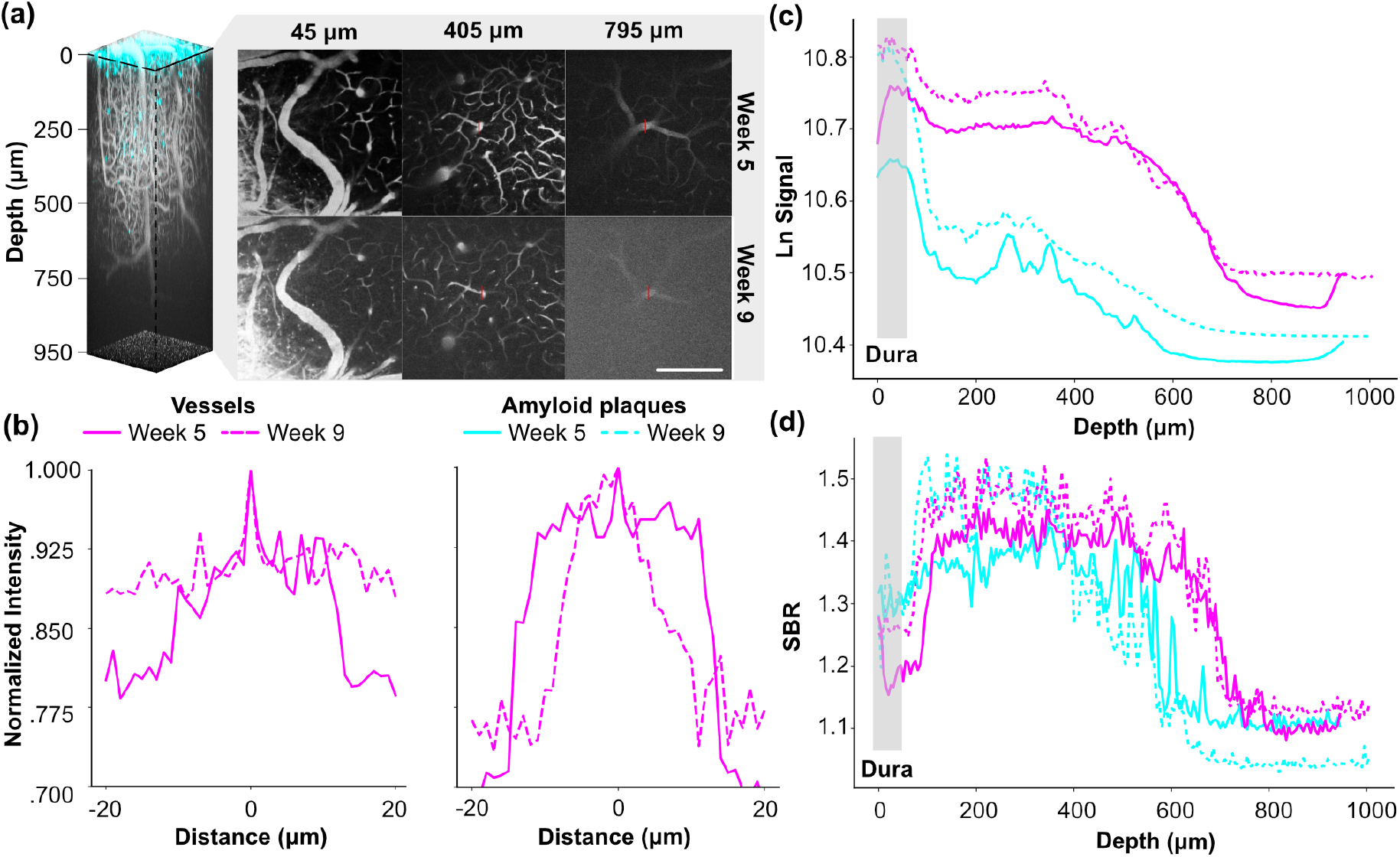
Signal-to-background ratio (SBR) and signal strength analysis over depth. (a) 3D reconstruction of vessels and plaque image stack. The amyloid plaques are labelled with methoxy-XO4 (blue), and the vessels are labelled with Texas Red (grey). The excitation wavelength is 1340 nm, the maximum laser power was 182 mW under the objective lens, and images were taken with 5 *µ*m z-steps. Shown are images taken with 25x objective lens at different depths (45, 405, 795 *µ*m). The brightest vessel at 405 and 795 *µ*m are marked in red. (b) Line profiles of the vessels highlighted with the brightest markers in red at depths 405 and 795 *µ*m. Scale bars is equal to 200 *µ*m. (c) The signal strength of the vessels was quantified by averaging the intensity of the top 0.5% of pixels in each frame.^32^ (d) The SBR was determined as the average of the brightest 15 and darkest 15 pixels in each profile. The SBR decreased with increasing depth, and there was an overall higher SBR in the first imaging session.^53^

To evaluate the reproducibility and depth-resolved imaging capabilities of our 3PM setup, we analyzed intensity profiles of the brightest vessels at depths of 405 and 795 *µ*m across both imaging sessions [Fig. 4(b)]. The brightest vessels were identified by averaging the intensity of the top 0.5% of pixels in each image slice. Normalized line profiles demonstrated consistent peak intensities at the same spatial locations between sessions, confirming reliable re-identification of vascular structures over time.

We quantified signal intensity and SBR for both vascular and amyloid signals as a function of imaging depth [Figs. 4(c) and 4(d)]. Vessel signals exhibited consistently higher SBR values compared to amyloid plaques at all depths, indicating superior contrast and detectability for vascular structures during deep imaging. Both signals displayed an initial increase near the cortical surface, attributed to higher tissue density in the dura mater, followed by a decline with depth due to light scattering and absorption. The dura mater location was determined using the THG signal at the top of the stack. Variations in signal strength and SBR between sessions were primarily attributable to differences in laser power settings and dye concentrations (Texas Red for vessels and methoxy-XO4 for amyloid plaques). To minimize these effects, raw data were processed using adaptive histogram normalization, and the dura mater was excluded during segmentation to avoid artifacts.

### 3.3 Overall vessel volume decreased and amyloid plaques increased in the second session

After segmentation, vascular and amyloid volumes were quantified across two imaging sessions in four longitudinally matched FOVs from three mice aged 6.5 to 9.9 months. Percent change in vessel volume between sessions (S2 vs. S1) ranged from –36.6% to 4.3%, while percent change in plaque volume ranged from 48.2% to 730.2%. The data shows a vessel volume decrease in 3 FOVs, while the plaque volume increased in all FOVs.

### 3.4 Significant Vessel and Plaque Changes Across Sessions

As shown in Figure 6, vessel diameter decreased significantly between the two imaging sessions for three of the four FOVs (Mann–Whitney U tests: all *p* < 0.001). Vessel length significantly increased for Mouse 1 FOV 1 (*p* < 0.001) and decreased FOV 2 (*p* = 0.031), whereas no significant difference was observed for Mouse 2 FOV 2 or Mouse 3 FOV 1 (*p* > 0.05). Vessel tortuosity did not differ significantly between sessions for any FOV (all *p* > 0.05). Inter-vessel distance increased in the second session for both Mouse 1 FOV 1 and Mouse 2 FOV 2 (both *p* < 0.001). Plaque radius significanlty increased only for Mouse 1 FOV 2 (*p* < 0.05), with no other significant differences observed across FOVs (*p* > 0.05).

## 4 Discussion

In this proof-of-concept study, we established a longitudinal three-photon microscopy (3PM) approach for imaging vascular and amyloid pathologies in deep cortical structures of of APP^NL-G-F^ knock-in mice, a model of AD. This methodology overcomes the depth limitations of two-photon microscopy (2PM) and enables in vivo imaging of cortical structures at high resolution over extended periods. Although the present dataset is limited in size (four FOVs in three animals) and duration, it demonstrates the feasibility of using 3PM to monitor structural changes associated with AD pathology over time with increased sample size.

### 4.1 Longitudinal Three-Photon Imaging of Cortical Structures

Our imaging methodology enabled longitudinal visualization of amyloid plaques at cortical depths of up to 420 *µ*m, together with vascular structures observable to depths of up to 920 *µ*m [Fig. 3(a)], which exceeds the previously reported range of 150 to 200 *µ*m achieved in *in vivo* studies of amyloid plaques using two-photon microscopy (2PM).^30, 37^ While longitudinal studies using MRI and PET have achieved greater imaging depths,^54, 55^ they do so at a significantly lower resolution. 3PM, with its cellular resolution, bridges the gap between superficial multiphoton imaging and macroscopic modalities like MRI and PET, and has the capability to analyze neuronal circuit activity in addition to amyloid plaque load and vasculature.

In this paper, we achieved a maximum imaging depth of 1.09 mm [Fig. 3(b)] in a 5-week old mouse, which is consistent with the previously reported depth of 1.1 mm by Ouzounov et al.,^31^ using the same excitation wavelength of 1340 nm. In older animals (6.5–9.9 months), the attainable imaging depth was reduced, likely caused by increased light scattering in more mature brain tissue, limiting optical penetration.^56^. Improvements can possibly be expected with the use of longer wavelengths or optimized immersion media such as deuterium oxide (D_2_O).^36^ Imaging of AAV-mediated GFP expression alongside vascular labeling confirmed spectral compatibility and minimal cross-talk at 1340 nm, supporting the feasibility of multicolor 3PM for simultaneous imaging of genetically and chemically labeled structures.

Field-of-view consistency across imaging sessions was ensured using vascular landmarks and stable mounting techniques [Fig. 4(a)], with normalized line profiles demonstrating reproducibility [Fig. 4(b)]. This highlights the feasibility of 3PM for longitudinal studies, offering high resolution with minimal photodamage due to longer excitation wavelengths and low average laser power.^33^ The combined use of Texas Red–dextran and methoxy-X04 for simultaneous in vivo visualization of cerebral vasculature and amyloid plaques is well established and has been validated in prior multiphoton imaging studies.^57^ However, future studies aiming to further integrate structural and molecular readouts could incorporate post-imaging immunohistochemical staining. CD31 for endothelial cells, and Thioflavin-S or 6E10 for amyloid plaques could be used to compare longitudinal *in vivo* dynamics with fixed-tissue histopathology, thereby deepening the biological interpretation of three-photon datasets.^58, 59^

### 4.2 Vascular Degeneration and Amyloid Accumulation

Although biological interpretations cannot be drawn from this limited dataset, the longitudinal imaging pipeline captured significant variations in vascular and amyloid volumes over time [Fig. 5–6]. Prior studies in AD models have consistently reported reductions in vessel density and diameter, increased inter-vessel spacing, and general vascular rarefaction accompanying amyloid accumulation.^60–71^

**Fig 5.**
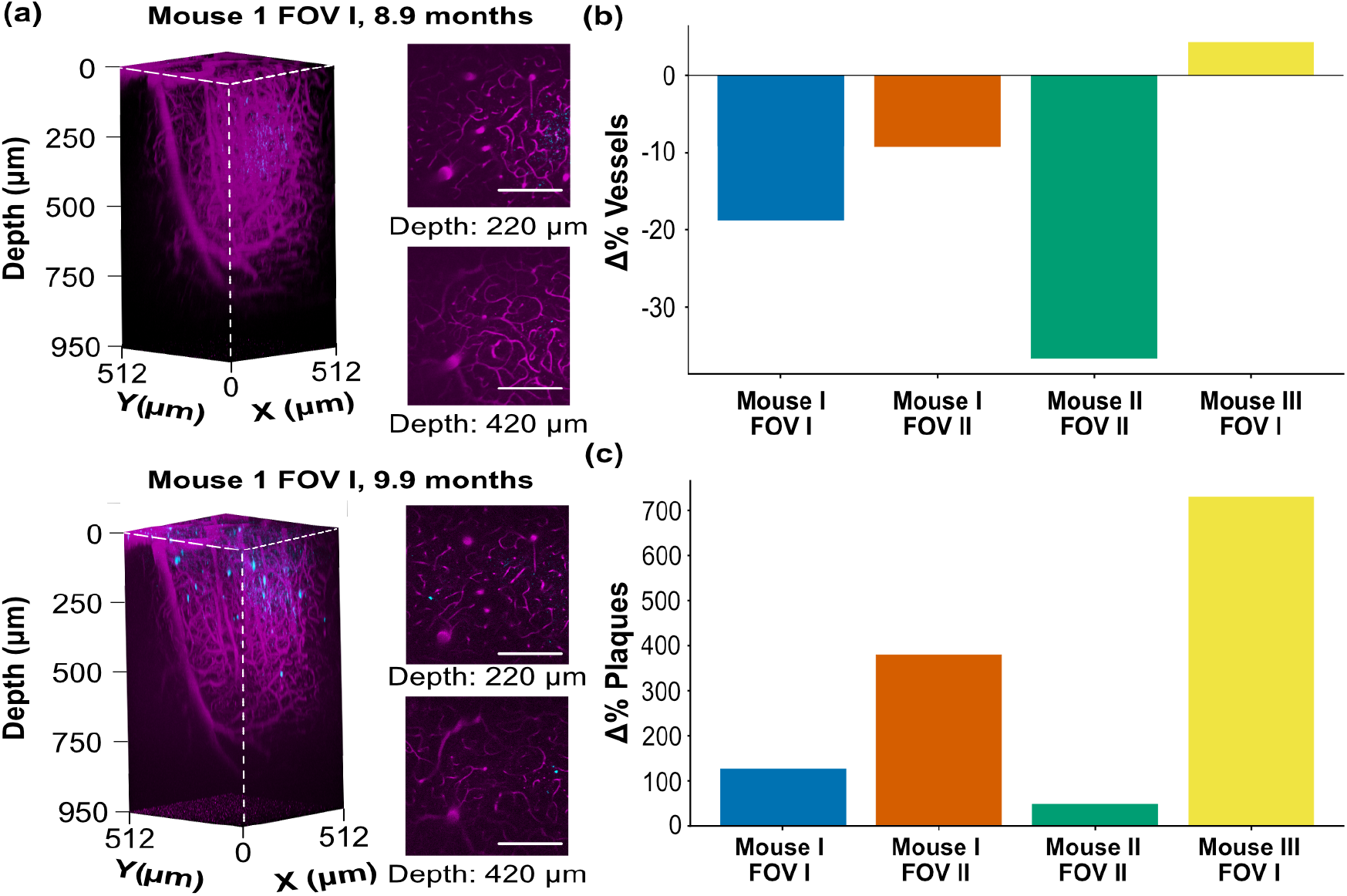
Vessel and amyloid plaques volume changes between imaging session 1 and 2. (a) Representative three-photon fluorescence renderings of vasculature (magenta) and amyloid plaques (cyan) from Mouse 1 FOV I at Session 1 (top) and Session 2 (bottom), with example slices shown at 220 µm and 420 µm depth. (b) Percent change in vessel volume (S2 vs. S1) across four longitudinally matched FOVs from three mice. (c) Percent change in plaque volume (S2 vs. S1). Each bar represents one FOV. Scale bars are 200 *µ*m.

**Fig 6.**
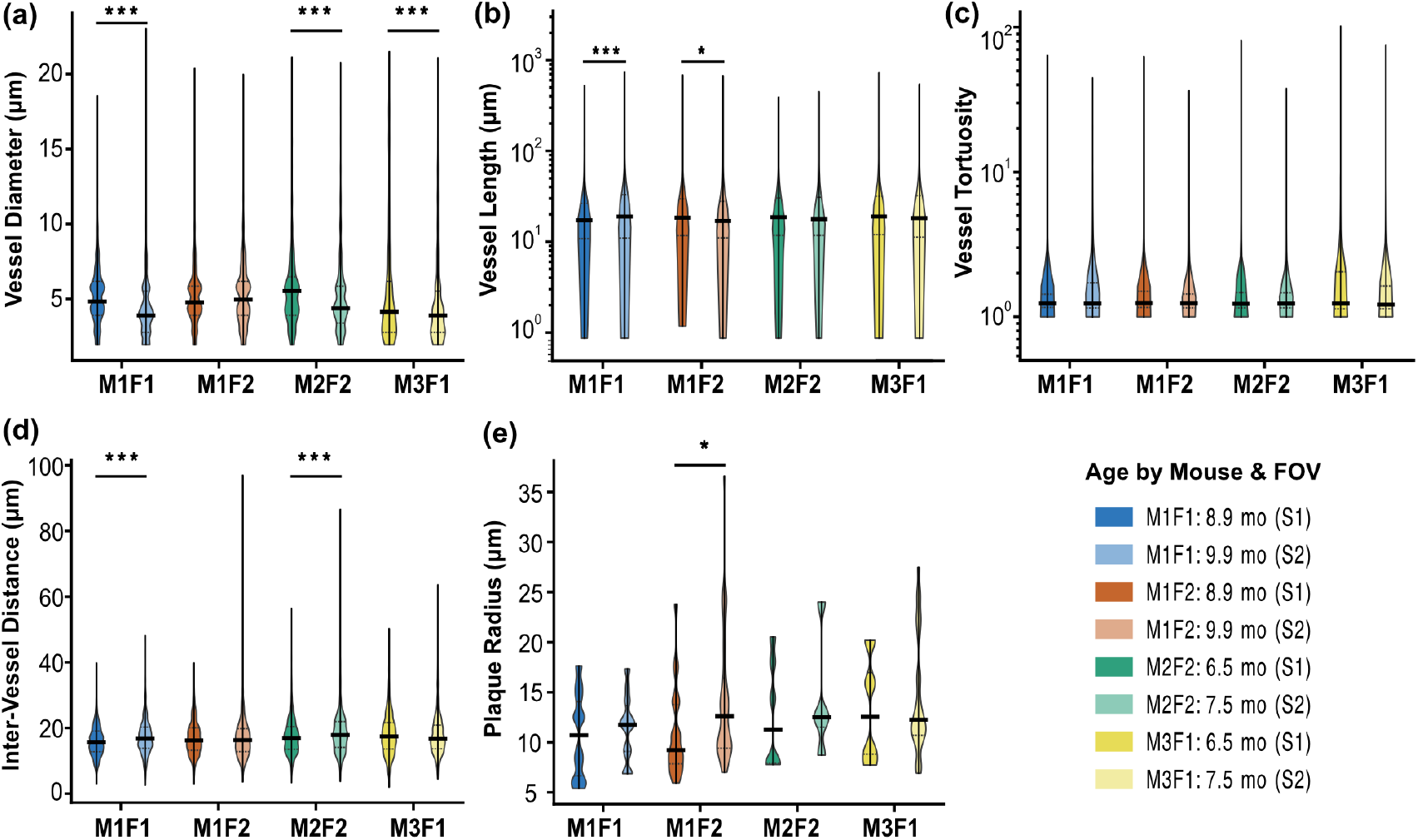
Change of vascular parameters between imaging session 1 and 2. Subplots show (a) vessel diameter, (b) vessel length, (c) vessel tortuosity, (d) inter-vessel distance, and (e) plaque radius. Data was tested for normalization and statistical analysis was done using the Mann-Whitney *U* test. p values are indicated by stars, * p<0.05, ** p<0.01 and *** p<0.001.

It should be noted that apparent vascular changes could be influenced by technical or physiological factors such as minor compression during imaging, tissue swelling, or variability in vascular dye perfusion. Furthermore, the four-week imaging interval may be insufficient to capture slow vascular remodeling relative to amyloid deposition. The relatively subtle plaque-associated effects are consistent with the slower progression characteristic of the heterozygous NL-G-F genotype compared to more aggressive or homozygous AD models.^22, 23^

Overall, this pilot dataset demonstrates that 3PM can quantify vascular and amyloid structures in deep cortex across imaging sessions, establishing a foundation for future studies. Expanding this approach to larger sample sizes, multiple cortical regions, and longer monitoring intervals will allow quantitative assessment of the temporal relationship between vascular remodeling and amyloid accumulation in AD and possibly its response to therapeutic interventions.

## 5 Conclusion

We developed a longitudinal 3PM imaging methodology for studying vascular and amyloid pathologies in deep cortical structures of APP^NL-G-F^ mice. This approach could provides detailed insights into AD progression and surpasses traditional depth limitations of 2PM. Future research should leverage this methodology with larger cohorts, extended monitoring periods, and advanced analytical tools, such as machine learning algorithms, to further elucidate the interplay between vascular and amyloid pathologies in AD and evaluate potential therapeutic strategies.

## Code and Data Availability

Code used to generate the figures in this paper can be found on https://github.com/estassss/Longitudinal-three-photon-imaging. Data is available on https://doi.org/10.5061/dryad.wh70rxx2j.

## Ethics Statement

The animal study was reviewed and approved by Imperial College Animal Welfare Ethical Review Board and authorized under Home Office Project License with PPL number PP6988384.

## Author Contributions

ES, MY, MA and SS designed the experiments. ES and MY worked out the technical details of the experiments. MY performed the surgeries. ES and MG performed the experiments. ES analysed the data with support from MY, MG, and SS. ES wrote the manuscript. ES, MY, MA and SS supervised the project. All authors discussed the results and commented on the manuscript.

## Acknowledgments

We would like to thank Nawal Zabouri for her advice and insights on the surgical procedures used in the paper.

**Eline Stas** received her MSc degree in Engineering for Biomedicine from Imperial College London and was trained in three photon microscopy, image analysis and Alzheimer’s Disease in the lab of Neural Coding and Neurodegenerative Diseases of Prof. Simon Schultz.

**Mengke Yang** completed his PhD in Optics at the University of the Chinese Academy of Sciences. Following this, he conducted postdoctoral research at the Dementia Research Institute of University College London and in the Neural Coding and Neurodegenerative Diseases lab of Prof. Simon Schultz in the Department of Bioengineering at Imperial College London. His research focuses on ultra-large-field multiphoton imaging, related imaging techniques, and the study of neurodegenerative diseases such as Alzheimer’s disease.

**Simon Schultz** is a Professor of Neurotechnology and the Director of the Centre for Neurotechnology at Imperial College London. He began his academic career studying physics, applied mathematics, and electrical engineering at Monash University and the University of Sydney. He then pursued a DPhil in computational neuroscience at the University of Oxford. Following his doctorate, he conducted postdoctoral research in experimental neuroscience at New York University with Tony Movshon and at University College London with Michael Haüsser. In 2004, he joined Imperial College London, where he continues to make significant contributions to the field of neurotechnology.

**Mary Ann Go** trained in physics before undertaking a PhD in Neuroscience at the Australian National University. She then conducted postdoctoral research at Imperial College London in the Neural Coding and Neurodegenerative Diseases lab of Prof. Simon Schultz. Her research interests involve the development and use of optics/photonics technologies to study the nervous system and neurodegenerative diseases such as Alzheimer’s disease.

---

**Fig S1.**
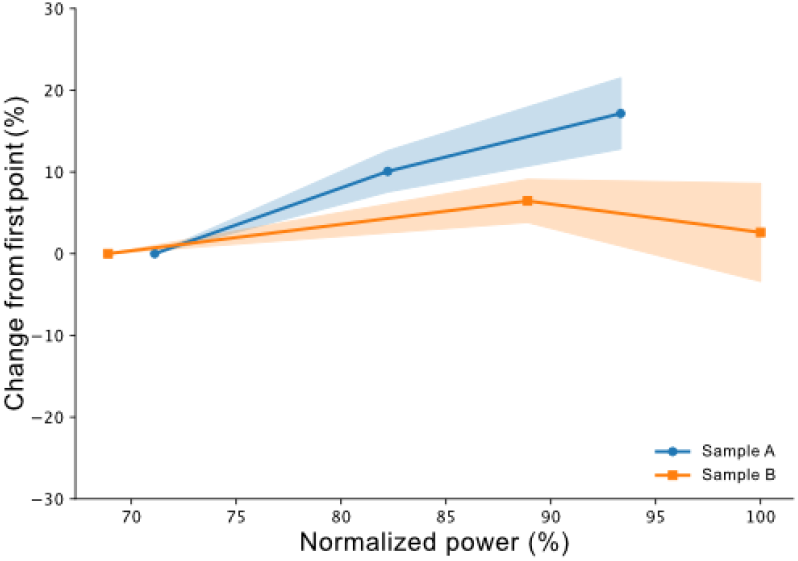
Stability of measurement. Measured width of structures in a neonatal mouse lung slice imaged under varying power levels. (b) Shift of selected vessels (relative to previous slice) per depth of FOV 1 Session 1 for the three mice.

**Fig S2.**
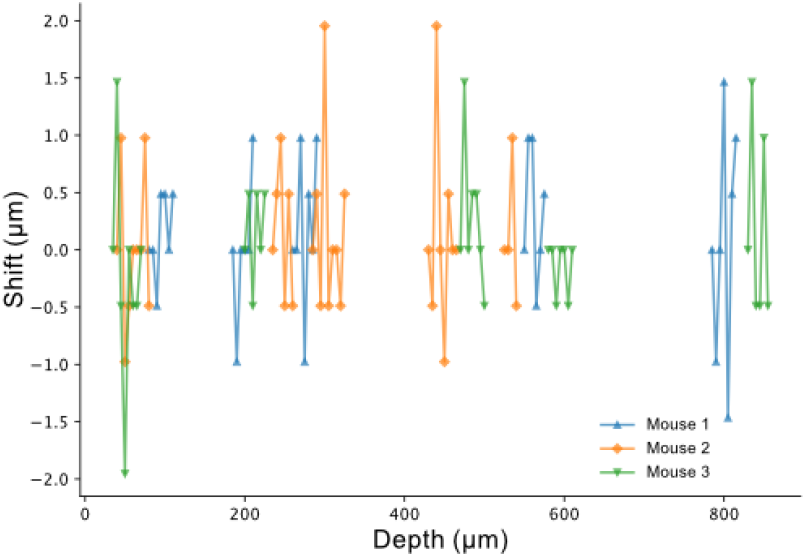
Motion artifact assessment. Shift of selected vessels (relative to previous slice) per depth of FOV 1 Session 1 for the three mice.

## References

1 H. Hampel, J. Hardy, K. Blennow, et al., “The amyloid-β pathway in alzheimer’s disease,” Molecular Psychiatry 26, 5481–5503 (2021).

2 J. A. Hardy and G. A. Higgins, “Alzheimer’s disease: The amyloid cascade hypothesis,” Science 256(5054), 184–186 (1992).

3 G. S. Bloom, “Amyloid- and tau: the trigger and bullet in alzheimer disease pathogenesis,” JAMA Neurology 71(4), 505–508 (2014).

4 E. J. Solis, K. N. Hascup, and E. R. Hascup, “Alzheimer’s disease: The link between amyloid- and neurovascular dysfunction,” Journal of Alzheimer’s Disease 76(4), 1179–1198 (2020).

5 J. N. Rauch and L. C. Hsieh-Wilson, “Mechanisms of pathogenic tau and a protein spreading in alzheimer’s disease,” Frontiers in Aging Neuroscience 12(265) (2020).

6 J. Klohs, “An integrated view on vascular dysfunction in alzheimer’s disease,” Neurodegenerative Diseases 19, 109–127 (2020).

7 Y. Take, Y. Chikai, K. Shimamori, et al., “Amyloid aggregation induces human brain microvascular endothelial cell death with abnormal actin organization,” Biochemical and Biophysical Reports 29, 101189 (2022).

8 J. Li, M. Li, Y. Ge, et al., “-amyloid protein induces mitophagy-dependent ferroptosis through the cd36/pink/parkin pathway leading to blood-brain barrier destruction in alzheimer’s disease,” Cell & Bioscience 12, 69 (2022).

9 X. Sun, G. He, H. Qing, et al., “Hypoxia facilitates alzheimer’s disease pathogenesis by up-regulating bace1 gene expression,” Proceedings of the National Academy of Sciences 103(49), 18727–18732 (2006).

10 R. Nortley, N. Korte, P. Izquierdo, et al., “Amyloid oligomers constrict human capillaries in alzheimer’s disease via signaling to pericytes,” Science 365(6450), eaav9518 (2019).

11 V. Fischer, A. Siddiqi, and Y. Yusufaly, “Altered angioarchitecture in selected areas of brains with alzheimer’s disease,” Acta Neuropathologica 79, 672–679 (1990).

12 M. D. Sweeney, A. Montagne, A. P. Sagare, et al., “Vascular dysfunction—the disregarded partner of alzheimer’s disease,” Alzheimers Dement. 15, 158–167 (2019).

13 J. Beskow, O. Hassler, and J. Ottosson, “Cerebral arterial deformities in relation to senile deterioration,” Acta Psychiatrica Scandinavica 47, 111–119 (1971).

14 V. R. Challa, C. R. Thore, D. M. Moody, et al., “Increase of white matter string vessels in Alzheimer’s disease,” Journal of Alzheimer’s Disease: JAD 6(4), 379–449 (2004).

15 N. Wang, X. Yang, Z. Zhao, et al., “Cooperation between neurovascular dysfunction and a in alzheimer’s disease,” Front. Mol. Neurosci. 16, Article 1227493 (2023).

16 S. A. Kozin, O. I. Kechko, A. A. Adzhubei, et al., “Switching on/off amyloid plaque formation in transgenic animal models of alzheimer’s disease,” International Journal of Molecular Sciences 25(1), 72 (2024).

17 M. Yokoyama, H. Kobayashi, L. Tatsumi, et al., “Mouse models of alzheimer’s disease,” Frontiers in Molecular Neuroscience 15, 912995 (2022).

18 A. Latif-Hernandez, V. Sabanov, T. Ahmed, et al., “The two faces of synaptic failure in appNL−G−F knock-in mice,” Alzheimer’s Research & Therapy 12, 100 (2020).

19 T. Saito, Y. Matsuba, N. Mihira, et al., “Single app knock-in mouse models of alzheimer’s disease,” Nature Neuroscience 17, 661–663 (2014).

20 H. Sasaguri, P. Nilsson, S. Hashimoto, et al., “App mouse models for alzheimer’s disease preclinical studies,” EMBO Journal 36, 2473–2487 (2017).

21 H. M. Lanoiseleé, G. Nicolas, D. Wallon, et al., “App, psen1, and psen2 mutations in earlyonset alzheimer disease: A genetic screening study of familial and sporadic cases,” PLOS Medicine 14, e1002270 (2017).

22 S. E. B. Maezono, M. Kanuka, C. Tatsuzawa, et al., “Progressive changes in sleep and its relations to amyloid- distribution and learning in single app knock-in mice,” eNeuro 7 (2020).

23 D. Kwart, A. Gregg, C. Scheckel, et al., “A large panel of isogenic app and psen1 mutant human ipsc neurons reveals shared endosomal abnormalities mediated by app-ctfs, not a,” Neuron 104, 256–270.e5 (2019).

24 S. Heinzer, T. Krucker, M. Stampanoni, et al., “Hierarchical bioimaging and quantification of vasculature in disease models using corrosion casts and microcomputed tomography,” in Proc. SPIE, 5535, 65–76, SPIE (2004).

25 M. Haft-Javaherian, L. Fang, V. Muse, et al., “Deep convolutional neural networks for segmenting 3d in vivo multiphoton images of vasculature in alzheimer disease mouse models,” PLoS ONE 14(3), e0213539 (2019).

26 S. J. Moore et al., “Degeneration of vascular architecture associated with disease pathology in the 5xfad mouse model of alzheimer’s disease,” 16(S1), e041262 (2020).

27 G. R. Frost, V. Longo, T. Li, et al., “Hybrid pet/mri enables high-spatial resolution, quantitative imaging of amyloid plaques in an alzheimer’s disease mouse model,” Scientific Reports 10, Article number 10379 (2020).

28 M. Markicevic, I. Savvateev, C. Grimm, et al., “Emerging imaging methods to study wholebrain function in rodent models,” Translational Psychiatry 11(457) (2021).

29 A. R. Kherlopian, T. Song, Q. Duan, et al., “A review of imaging techniques for systems biology,” BMC Systems Biology 2(1), 1–18 (2008).

30 J. K. Hefendehl, B. M. Wegenast-Braun, C. Liebig, et al., “Long-term in vivo imaging of -amyloid plaque appearance and growth in a mouse model of cerebral-amyloidosis,” The Journal of Neuroscience 31(2), 624–629 (2011).

31 D. G. Ouzounov, T. Wang, M. Wang, et al., “In vivo three-photon imaging of activity of gcamp6-labeled neurons deep in intact mouse brain,” Nature Methods 14(4), 388–390 (2017).

32 T. Wang, D. G. Ouzounov, C. Wu, et al., “Three-photon imaging of mouse brain structure and function through the intact skull,” Nature Methods 15(10), 789–792 (2018).

33 T. Wang, C. Wu, D. G. Ouzounov, et al., “Quantitative analysis of 1300-nm three-photon calcium imaging in the mouse brain,” eLife 9, e53205 (2020).

34 Y. Xiao, P. Deng, Y. Zhao, et al., “Three-photon excited fluorescence imaging in neuroscience: From principles to applications,” Frontiers in Neuroscience 17, 1–21 (2023).

35 N. G. Horton, K. Wang, D. Kobat, et al., “In vivo three-photon microscopy of subcortical structures within an intact mouse brain,” Nature Photonics 7, 205–209 (2013).

36 Y. Hontani, F. Xia, and C. Xu, “Multicolor three-photon fluorescence imaging with single-wavelength excitation deep in mouse brain,” Science Advances 7(eabf3531) (2021).

37 Z. Luo, H. Xu, S. Samanta, et al., “Long-term repeatable in vivo monitoring of amyloid-plaques and vessels in alzheimer’s disease mouse model with combined tpef/cars microscopy,” Biomedicines 10, 2949 (2022).

38 L. P. Munting, M. P. P. Derieppe, L. M. Voortman, et al., “Multi-scale assessment of brain blood volume and perfusion in the app/ps1 mouse model of amyloidosis,” bioRxiv (2022).

39 M. A. Go, J. Rogers, G. P. Gava, et al., “Place cells in head-fixed mice navigating a floating real-world environment,” Frontiers in Cellular Neuroscience 15, 618658 (2021).

40 A. Holtmaat, T. Bonhoeffer, D. K. Chow, et al., “Long-term, high-resolution imaging in the mouse neocortex through a chronic cranial window,” Nature Protocols 4(8), 1128–1144 (2009).

41 S. E. Gombash Lampe, B. K. Kasper, and K. D. Foust, “Intravenous injection in neonatal mice,” J. Vis Exp. 11(93), e52037 (2014).

42 T. A. Pologruto, B. L. Sabatini, and K. Svoboda, “Scanimage: Flexible software for operating laser scanning microscopes,” Biomedical Engineering OnLine 2(13) (2003).

43 C.-L. Chiu, N. Clack, and the napari community, “napari: a python multi-dimensional image viewer platform for the research community,” Microscopy and Microanalysis 28(S1), 1576–1577 (2022).

44 P. Spangenberg, N. Hagemann, A. Squire, et al., “Rapid and fully automated blood vasculature analysis in 3d light-sheet image volumes of different organs,” Cell Rep Methods 3(3), 100436 (2023).

45 C. Kirst, S. Skribiane, A. Vieites-Prado, et al., “Mapping the fine-scale organization and plasticity of the brain vasculature,” Cell 180(4), 780–795.e25 (2020).

46 C. L. Walsh, M. Berg, H. West, et al., “Reconstructing microvascular network skeletons from 3d images: What is the ground truth?,” Computers in Biology and Medicine 171, 108140 (2024).

47 B. Garcia-Garcia, H. Mattern, N. Vockert, et al., “Vessel distance mapping: A novel methodology for assessing vascular-induced cognitive resilience,” NeuroImage 274, 120094 (2023).

48 P. Virtanen, R. Gommers, T. E. Oliphant, et al., “Scipy: Open source scientific tools for python,” (2024).

49 N. Watanabe, Y. Noda, T. Nemoto, et al., “Cerebral artery dilation during transient ischemia is impaired by amyloid deposition around the cerebral artery in alzheimer’s disease model mice,” The Journal of Physiological Sciences 70(57) (2020).

50 M. Kavkova, T. Zikmund, A. Kala, et al., “Contrast enhanced x-ray computed tomography imaging of amyloid plaques in alzheimer disease rat model on lab based micro ct system,” Scientific Reports 11, Article number: 5999 (2021).

51 P. Virtanen, R. Gommers, T. E. Oliphant, et al., “SciPy 1.0: Fundamental Algorithms for Scientific Computing in Python,” Nature Methods 17, 261–272 (2020).

52 F. Pedregosa, G. Varoquaux, A. Gramfort, et al., “Scikit-learn: Machine learning in python,” Journal of Machine Learning Research 12, 2825–2830 (2011).

53 S. A. Engelmann, A. Tomar, A. L. Woods, et al., “Pulse train gating to improve signal generation for in vivo two-photon fluorescence microscopy,” bioRxiv (2023).

54 J. Klohs, I. W. Politano, A. Deistung, et al., “Longitudinal assessment of amyloid pathology in transgenic arca mice using multi-parametric magnetic resonance imaging,” PLOS ONE 8(6), e66097 (2013).

55 J. Maeda, B. Ji, T. Irie, et al., “Longitudinal, quantitative assessment of amyloid, neuroinflammation, and anti-amyloid treatment in a living mouse model of alzheimer’s disease enabled by positron emission tomography,” The Journal of Neuroscience 27(41), 10957–10968 (2007).

56 M. Oheim, E. Beaurepaire, E. Chaigneau, et al., “Two-photon microscopy in brain tissue: parameters influencing the imaging depth,” 111(1), 29–37 (2001).

57 M. Meyer-Luehmann, T. L. Spires-Jones, C. Prada, et al., “Rapid appearance and local toxicity of amyloid-β plaques in a mouse model of alzheimer’s disease,” Nature 451, 720–724 (2008).

58 M. P. Pusztaszeri, W. Seelentag, and F. T. Bosman, “Immunohistochemical expression of endothelial markers cd31, cd34, von willebrand factor, and fli-1 in normal human tissues,” Journal of Histochemistry & Cytochemistry 54(4), 385–395 (2006).

59 P. Giannoni, M. Arango-Lievano, I. Das Neves, et al., “Cerebrovascular pathology during the progression of experimental alzheimer’s disease,” Neurobiology of Disease 88, 107–117 (2016).

60 H. Oakley, S. L. Cole, S. Logan, et al., “Intraneuronal-amyloid aggregates, neurodegeneration and neuron loss in transgenic mice with five familial Alzheimer’s disease mutations: potential factors in amyloid plaque formation,” Journal of Neuroscience 26, 10129–10140 (2006).

61 A. Jullienne, J. I. Szu, R. Quan, et al., “Cortical cerebrovascular and metabolic perturbations in the 5xFAD mouse model of Alzheimer’s disease,” 15, 1220036 (2023).

62 S. Dudal, P. Krzywkowski, J. Paquette, et al., “Inflammation occurs early during the a deposition process in tgcrnd8 mice,” Neurobiology of Aging 25, 861–871 (2004).

63 A. Dorr, B. Sahota, L. V. Chinta, et al., “Amyloid--dependent compromise of microvascular structure and function in a model of Alzheimer’s disease,” Brain 135, 3039–3050 (2012).

64 D. Van Dam, R. D’Hooge, M. Staufenbiel, et al., “Age-dependent cognitive decline in the APP23 model precedes amyloid deposition,” European Journal of Neuroscience 17, 388–396 (2003).

65 E. P. Meyer, A. Ulmann-Schuler, M. Staufenbiel, et al., “Altered morphology and 3d architecture of brain vasculature in a mouse model for Alzheimer’s disease,” Proceedings of the National Academy of Sciences of the United States of America (PNAS) 105, 3587–3592 (2008). Freely available online through the PNAS open access option.

66 W. Kamphuis, C. Mamber, M. Moeton, et al., “GFAP isoforms in adult mouse brain with a focus on neurogenic astrocytes and reactive astrogliosis in mouse models of Alzheimer’s disease,” PLOS ONE 7(e42823) (2012).

67 G. D. Lee, J. H. Aruna, P. M. Barrett, et al., “Stereological analysis of microvascular parameters in a double transgenic model of Alzheimer’s disease,” 65, 317–322 (2005).

68 D. Games, D. Adams, R. Alessandrini, et al., “Alzheimer-type neuropathology in transgenic mice overexpressing v717f-amyloid precursor protein,” Nature 373, 523–527 (1995).

69 C. Pomilio, J. Presa, C. Oses, et al., “Loss of direct vascular contact to astrocytes in the hippocampus as an initial event in Alzheimer’s disease: Evidence from patients, in vivo and in vitro experimental models,” Molecular Neurobiology 61, 5142–5160 (2024).

70 S. A. Frautschy, F. Yang, M. Irrizarry, et al., “Microglial response to amyloid plaques in APPsw transgenic mice,” American Journal of Pathology 152, 307–317 (1998).

71 Y. Zhang, F. L. Chao, L. Zhang, et al., “Quantitative study of the capillaries within the white matter of the tg2576 mouse model of Alzheimer’s disease,” 9(4), e01268 (2019).

